# bioq: a unified, agent-native command-line interface to a fleet of AI drug-discovery methods

**DOI:** 10.64898/2026.09.03.749148

**Authors:** Zhiheng Liu, Yinan Wang

## Abstract

Artificial intelligence methods now span the drug-discovery pipeline—structure prediction, de novo design, docking, affinity estimation, and ADMET—yet each tool ships with its own, often incompatible, GPU software stack, so chaining several into a workflow requires reproducing conflicting software environments and access to datacenter-class hardware that most labs do not have. **bioq** is a dependency-light command-line client backed by **bioq-services** (a control-plane gateway plus a growing fleet of 38+ containerized drug-discovery tools spanning 7 discovery stages and 6 molecular modalities). The bioq CLI is self-describing and consistent across the fleet of computational tools, giving researchers and coding agents uniform access to any tool from a laptop, with no local model code, CUDA setup, or cloud configuration, running on serverless GPUs billed per job. The interface–gateway–services architecture makes bioq an execution substrate for automated, agent-driven discovery. bioq and bioq-services are open source under the MIT License and available at https://github.com/wolfsonliu/bioq and https://github.com/wolfsonliu/bioq-services. bioq runs on Python ≥ 3.10 with `httpx` as its only runtime dependency; services run as Linux containers and are self-hostable via the provided local deployment scripts (alongside scripts for Alibaba Cloud Function Compute). Install, quickstart, test data, and self-hosting instructions are in the repository. The project is actively maintained and will continue to receive updates to both features and services.

## 1 Introduction

The past five years have seen a surge of AI methods across the drug-discovery pipeline, including structure prediction (AlphaFold (Jumper et al., 2021; Abramson et al., 2024), RoseTTAFold (Baek et al., 2021), Boltz (Wohlwend et al., 2024)), de novo protein design (RFdiffusion (Watson et al., 2023), ProteinMPNN (Dauparas et al., 2022)), small-molecule generation (REINVENT 4 (Loeffler et al., 2024)), molecular docking (DiffDock (Corso et al., 2023)), and affinity estimation (BindFlow (León et al., 2026)). Despite this progress, integrating these methods into a practical discovery workflow remains disproportionately difficult. The primary obstacle is often not the underlying science, but deployment: each tool typically depends on a specific combination of CUDA, conda packages, and deep-learning framework versions. These dependencies frequently conflict, making some tools impossible to install in a shared software environment (§4.1). Beyond software compatibility, computational accessibility presents a second fundamental barrier. Many of these deep-learning models require more GPU memory than is available on a laptop or typical workstation (§4.1), while purchasing and maintaining data-center-class GPUs is neither affordable nor practical for occasional use.

The life sciences research community has already tamed this kind of fragmentation through per-tool isolation: Bioconda (The Bioconda Team et al., 2018) for versioned packaging, BioContainers (Da Veiga Leprevost et al., 2017) for per-tool containers, and platforms such as Galaxy (The Galaxy Community et al., 2026) and nf-core (Langer et al., 2025), which together make analyses accessible and reproducible (Grüning et al., 2018). `bioq` adopts this one-container-per-tool model and extends it to meet the additional demands of modern AI-enabled discovery. In particular, many GPU-intensive models require hardware beyond the reach of a typical laboratory, yet they must still be composed into coherent workflows and exposed through a uniform interface that can be operated by both bench scientists and autonomous coding agents. ColabFold (Mirdita et al., 2022) demonstrated the value of making such models available without requiring users to own or maintain the underlying infrastructure. `bioq` generalizes this paradigm from a single model to an integrated discovery stack, providing a uniform interface and an architecture that can be readily extended with new tools.

Autonomous coding agents (such as Claude Code, Codex, opencode, DeepSeek Harness) further amplify this need. These agents can plan and execute computational research tasks and are increasingly being used to support scientific exploration (Gottweis et al., 2026; Ghareeb et al., 2026; Huang et al., 2026). Agents are particularly vulnerable to ecosystem fragmentation: limited access to local GPU hardware, numerous mutually incompatible software environments, and inconsistent tool-specific command-line interfaces can collectively slow down, or even derail, agent-driven research. A unified, dependency-light, self-describing interface provides precisely the abstraction that agents need; the same properties also lowers the barrier for a bench scientist. Our contributions are: (i) a uniform, self-describing `describe`/`run` contract for heterogeneous tools; (ii) dual-mode service images that support both HTTP serving and CLI-based batch execution, enabling reproducible runs across cloud and high-performance computing (HPC) environments; (iii) serverless GPU dispatch, eliminating the need for individual users to own or manage GPU infrastructure; and (iv) a curated fleet spanning the discovery stack, with a simple extension model in which each new tool requires only one container and one registry entry (§3).

## 2 Methods

### 2.1 Architecture

`bioq` has three tiers (Figure 1): the `bioq` thin client, a control-plane `gateway`, and a fleet of worker `services`. The gateway centralizes all platform-level functionality, including authentication, input and output storage, job dispatch, and service discovery through a registry. Authentication is based on OIDC/JWT, with identities managed by an external provider and jobs isolated by account. The client contains none of this platform logic.

**Figure 1.**
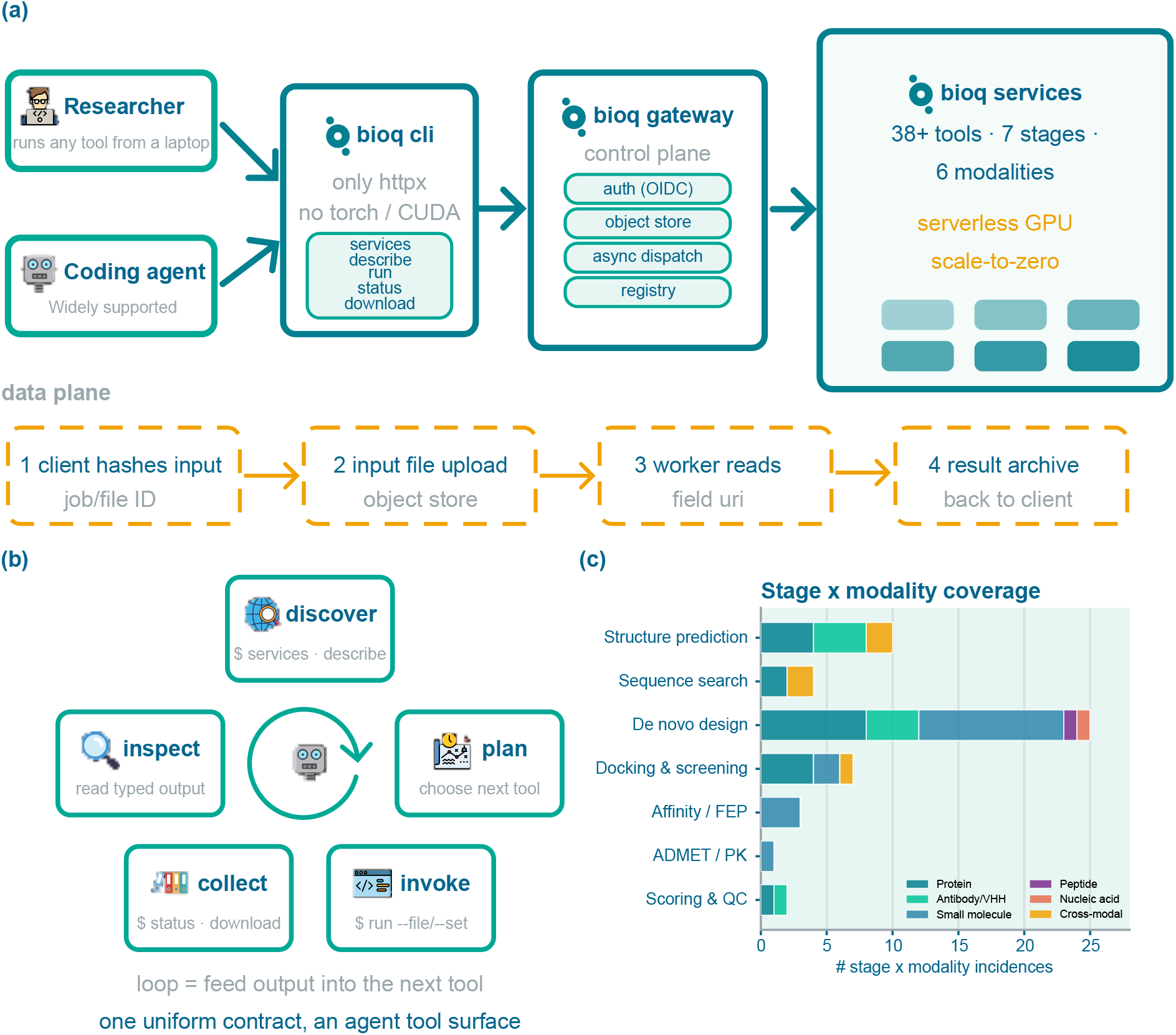
`bioq` unifies a three-tier service architecture behind a common interface for researchers and coding agents. (a) The thin `bioq` client communicates with a control-plane gateway, which dispatches jobs to more than 38 serverless GPU and CPU services. Large inputs are transferred through content-addressed storage and bypass the request path. (b) Coding agents use the same `describe`/`run` contract to form a closed loop of tool discovery, invocation, result collection, inspection, and chaining. (c) Modality coverage by discovery stage, with one stacked horizontal bar per stage.

A user or agent runs the `bioq` client on a local laptop, and the client communicates exclusively with the gateway over HTTPS. The gateway, in turn, dispatches jobs to serverless GPU workers that the user never provisions nor manage. These workers may run on a managed serverless platform such as Alibaba Cloud Function Compute (FC) or equivalent AWS service, or on a self-managed cluster. Because the client has few runtime dependency, installing `bioq` on a laptop requires only a standard `pip`/`uv` command; no local model code, GPU driver, or tool-specific environment is needed.

This centralized architecture allows a single administrator to deploy and operate the gateway and its associated services for an entire research group. For small or trusted groups, gateway authentication can be disabled, allowing clients to connect with minimal configuration.

### 2.2 A uniform, self-describing contract

Every tool, regardless of its native interface or input format, is exposed through the same five subcommands: `bioq services` lists the available services; `bioq describe <service>` displays a service manifest; `bioq run <service> <endpoint>` invokes an endpoint; and `bioq status` and `bioq download` retrieve job status and results, respectively. Each manifest is machine-readable and specifies the endpoints exposed by the service, the name and type of every parameter, and any required file inputs.

Jobs are submitted using a uniform syntax:

~~~
bioq run <service> <endpoint> \
--file field=path --set k=v --wait -o out
~~~

The option `–output json` returns structured, machine-readable output containing the job identifier and status. The manifest serves both as human-readable documentation and as a schema from which an agent can construct a valid invocation. Consequently, users and agents need to learn only one interface to access the entire service fleet.

To assess fleet-wide adherence to this contract, we performed a manifest-only audit of 30 deployed services comprising 141 endpoints, including 68 asynchronous task endpoints invocable through `bioq run`. Of these task endpoints, 98.5% were fully conformant, and the median per-service conformity score was 1.0, demonstrating that the contract is implemented consistently across the fleet (Figure S1).

### 2.3 Data plane and dual-mode execution

To transfer the large files consumed and produced by these models, `bioq` uses a content-addressed data plane. The client computes the SHA-256 hash of each input, obtains a presigned object-storage URL from the gateway, and uploads the file directly to object storage. Large inputs, such as protein structures (PDB files) and multiple-sequence-alignment (MSA) files, therefore bypass the gateway’s request path. Objects are deduplicated by content hash, so an input already present in storage need not be uploaded again. The gateway then passes the corresponding object URI to the job as `field_uri`, and the resulting artifacts are returned as a single archive.

Each service image supports two execution modes: an asynchronous HTTP job runner for cloud and serverless deployment, and a one-shot CLI batch program invoked as `python -m server <endpoint>`. The same image and parameter schema can therefore be used across managed serverless platforms, self-managed clusters, HPC batch systems, and local compute nodes, enabling reproducible execution without separate deployment-specific implementations.

In the serverless mode, the gateway dispatches jobs to scale-to-zero GPU functions (Jonas et al., 2019). When deployed on Alibaba Cloud Function Compute (FC), worker instances are provisioned on demand, billed according to job resource usage, and released when idle. This model gives users access to GPU capacity without requiring them to purchase, provision, or pay for idle hardware.

### 2.4 Access modes

`bioq` exposes the same contract to all consumers, whether human or agent. Human users interact with it through the command line or shell scripts. Autonomous coding agents use the identical interface, with only a single client dependency to install: `services` and `describe` provide structured tool discovery; `run`, together with `–file`, `–set`, and `–output json`, provides programmatic invocation; and a submit → poll → download → inspect loop enables the output of one tool to be passed to the next (Figure 1b). Because the interface is uniform, self-describing, and machine-readable, agents can discover tools and compose multistep workflows without requiring a bespoke adapter for each service.

## 3 The service fleet

The current fleet comprises 38 services spanning seven stages of the discovery pipeline and six molecular modalities: proteins, antibodies/VHHs, small molecules, peptides, nucleic acids, and cross-modal systems (Figure 1c). Of these services, 30 are currently deployed on our private serverless infrastructure hosted on FC; these services constitute the set audited in §2.2.

The fleet covers a broad range of computational tasks. It includes AlphaFold, Boltz2, and ESMFold2 for protein structure prediction; MMseqs2 (Steinegger and Söding, 2017) for sequence search; RFdiffusion, ProteinMPNN, Genie3, and RFantibody for de novo design, with RFantibody using an internal RoseTTAFold-2 (RF2) model for structure prediction and scoring (Bennett et al., 2026); DiffDock, HADDOCK3 (Giulini et al., 2025), and LightDock for molecular docking; BindFlow and QligFEP for affinity and free-energy estimation; OpenADMET (Fraser et al., 2026) for ADMET prediction; and DockQ (Basu and Wallner, 2016) and DeepRank-Ab (Xu et al., 2025) for model scoring. The full service list and its per-tool citations are in Supplementary Table S1.

The fleet is extensible by design. Adding a tool requires only a container image that implements the shared service contract and a corresponding registry entry. No modification to the client is required, and the newly deployed tool becomes discoverable through `bioq services`. The same architecture also allows the fleet to track upstream development efficiently. Since each tool is isolated in its own container and described declaratively, incorporating a new upstream release (or an entirely new method) requires rebuilding only the affected image, without altering any other service. The numbers reported above therefore represent a snapshot of a fleet designed to grow and evolve alongside the field.

## 4 Results

Because `bioq` addresses accessibility and integration rather than model accuracy, we evaluate the platform’s coverage and operational characteristics without re-benchmarking the predictive performance of the underlying models, which has been established in their respective upstream studies. We organize the results around three practical questions: how fragmented the underlying computational ecosystem is; how effectively `bioq` shields end users from this fragmentation; and how bioq-enabled workflows perform in terms of scalability, cost, and real-world usability.

### 4.1 Access barriers

The diversity of the underlying software stacks makes it impractical to consolidate the fleet into a single environment. An audit of all 38 services identified 15 distinct PyTorch versions, ranging from 1.12 to 2.12, across CUDA 11 and 12 and Python 3.9–3.12. In combination, these dependencies form 23 distinct stack signatures. Among the 32 tools for which parseable upstream dependency specifications were available, pairwise comparison showed that only 9.9% of tool pairs (49 of 496) were fully compatible, 48.6% of tool pairs (241 of 496) were hard-incompatible: their declared version constraints were disjoint, so no shared environment could satisfy both (Figure S2). `bioq` hides this complexity behind a single ∼1.8 MB client whose only runtime dependency is `httpx`.

Hardware availability presents a separate barrier. Most services in the fleet cannot run on the hardware available in a typical researcher’s laptop. An audit of per-service hardware requirements showed that the GPU services generally require more memory than laptop-class devices can provide. Among the 26 GPU services with quantified minimum VRAM requirements, 24 require more than 4GB and 14 require more than 8GB (Figure S3). `bioq` therefore executes these services on remote, serverless GPUs rather than on the client device.

### 4.2 Scale and operating cost

For embarrassingly parallel screening and design workloads, serverless dispatch provides throughput beyond that of a single workstation without requiring users to orchestrate workers themselves. In batches of 50 independent jobs, the observed speedup relative to serial execution reached approximately 16.8*×* for PLIP, 10.6*×* for ProteinMPNN, and 10.4*×* for REINVENT, with peak concurrency reaching approximately 49 worker instances across the evaluated workloads (Figures S4 and S5).

Because worker instances scale to zero when idle, compute costs are incurred primarily during job execution rather than through continuous GPU provisioning. Based on the public serverless pricing schedule, our cost model estimates a weighted mean cost of approximately $0.18 per job and a duty-cycle break-even point of approximately 8.8% relative to an owned A100 GPU (Figure S6). Serverless execution is therefore economically favorable under the assumptions of this model for bursty workloads that would otherwise leave dedicated hardware idle for most of its lifetime.

Scaling on Function Compute (FC) is ultimately constrained by cold-start latency and platform concurrency quotas. These limits affect short jobs and large bursts in particular, although concurrency quotas can be adjusted when additional capacity is required.

### 4.3 Use case: a de novo antibody-design campaign

As an end-to-end demonstration, we executed the RFantibody de novo antibody-design pipeline entirely through `bioq`. The pipeline comprises RFdiffusion for backbone generation, Protein-MPNN for sequence design, and RoseTTAFold-2 (RF2) for structure prediction and interface scoring (Bennett et al., 2026). For each target, the complete workflow was launched from a laptop using three uniform `bioq run` commands, without a local GPU, model installation, or CUDA environment:

~~~
bioq run rfantibody rfdiffusion -- file target= target. pdb -- file framework = vhh. pdb \
-- set num_designs =1000 -- set hotspots =… -- wait -o s1
bioq run rfantibody proteinmpnn -- file input_quiver=s1 /1 _rfdiffusion. qv \
-- set seqs_per_struct =8 -- wait -o s1
bioq run rfantibody rf2 -- file input_quiver=s1 /2 _proteinmpnn. qv -- wait -o s1
~~~

Although each stage uses a different model, `bioq` exposes all three through the same execution contract. Across nine targets—seven VHHs and two scFvs—the campaign generated 1,000 candidate backbones per target and eight sequences per backbone, for a total of 72,000 designed sequences. All jobs were dispatched to the scale-to-zero GPU workers described in §4.2, with no local model code, GPU provisioning, or CUDA configuration.

We evaluated the designs using the in silico filters defined in the original RFantibody study: interface predicted aligned error (pAE < 10 Å) and design–model CDR root-mean-square deviation (RMSD) below 2 Å, where the latter measures self-consistency between the designed backbone and the RF2-predicted structure. Under the best-of-eight criterion used by the upstream method, per-target acceptance rates ranged from 2.4% to 70.6%, consistent with the broad in silico acceptance funnel reported in the original study. Table S2 reports the complete per-target funnel and a comparison with the published RFantibody results. Figure S7 visualizes the per-target funnel, while Figures S8 and S9 show the corresponding interface-pAE and CDR-RMSD distributions.

The same execution pattern can be extended across independently packaged services—for example, using RFdiffusion for backbone generation, Boltz for structure prediction, and DockQ for scoring—without changing the underlying contract. Importantly, the acceptance rates reported here are computational filtering rates, not experimental binding success rates. In the upstream study, the experimental hit rate was at most 2%, substantially lower than the in silico acceptance rate (Bennett et al., 2026).

## 5 Discussion

`bioq` shifts access to computational models from tool-specific local installations to a uniform, remotely executable service contract. This approach removes much of the software-compatibility and hardware-provisioning burden for users who need to combine multiple tools, but it also introduces several trade-offs. Serverless throughput is bounded by cold-start latency and platform concurrency limits, and its economic advantage depends strongly on workload duty cycle. Above the estimated break-even point, dedicated GPU infrastructure is likely to be more cost-effective than cloud platform serverless computation.

The service fleet is curated and centrally versioned. This design makes it possible to add new methods and track upstream releases by rebuilding individual containers without disrupting the remainder of the fleet. For users who depend on multiple tools, such isolation avoids the “incompatibility tax” associated with maintaining mutually conflicting software environments. However, formal policies for release versioning, deprecation, backward compatibility, and long-term reproducibility remain to be established.

The current data plane assumes access to cloud-compatible object storage. Although this dependency can be mitigated through self-hosted storage and the CLI batch mode for HPC environments, fully disconnected or tightly regulated deployments may require additional integration. Workflow chaining is also currently implemented through client-side scripts. A declarative workflow layer, including explicit provenance, caching, retries, and dependency management, is left for future work; the uniform, self-describing contract introduced here provides the foundation on which such a layer can be built.

## Data and code availability

bioq and bioq-services are open source under the MIT License and are available at https://github.com/wolfsonliu/bioq and https://github.com/wolfsonliu/bioq-services. bioq runs on Python ≥ 3.10 with `httpx` as its only runtime dependency; services run as Linux containers and are self-hostable via the provided local deployment scripts (alongside scripts for Alibaba Cloud Function Compute). Install, quickstart, test data, and self-hosting instructions are in the repository. The project is actively maintained and will continue to receive updates to both features and services. Supplementary tables and figures (Figures S1–S9, Tables S1–S2) are provided as a separate supplementary PDF and are also available at https://github.com/wolfsonliu/bioq-paper.

## Acknowledgments

The authors thank the developers and maintainers of the open-source methods and platforms underlying this work for releasing their code and model weights, including AlphaFold, RoseTTAFold-2, ColabFold, RFdiffusion, ProteinMPNN, RFantibody, and MMseqs2, among others. LLM tools were used to assist with coding and language polishing. All scientific claims, analyses, and generated text were reviewed and verified by the authors, who take full responsibility for the final manuscript.

