## Supplementary for "bioq: a unified, agent-native command-line interface to a fleet of AI drug-discovery methods"

### Supplementary Information

bioq: a unified, agent-native gateway to a fleet of AI drug-discovery tools

#### Supplementary figures

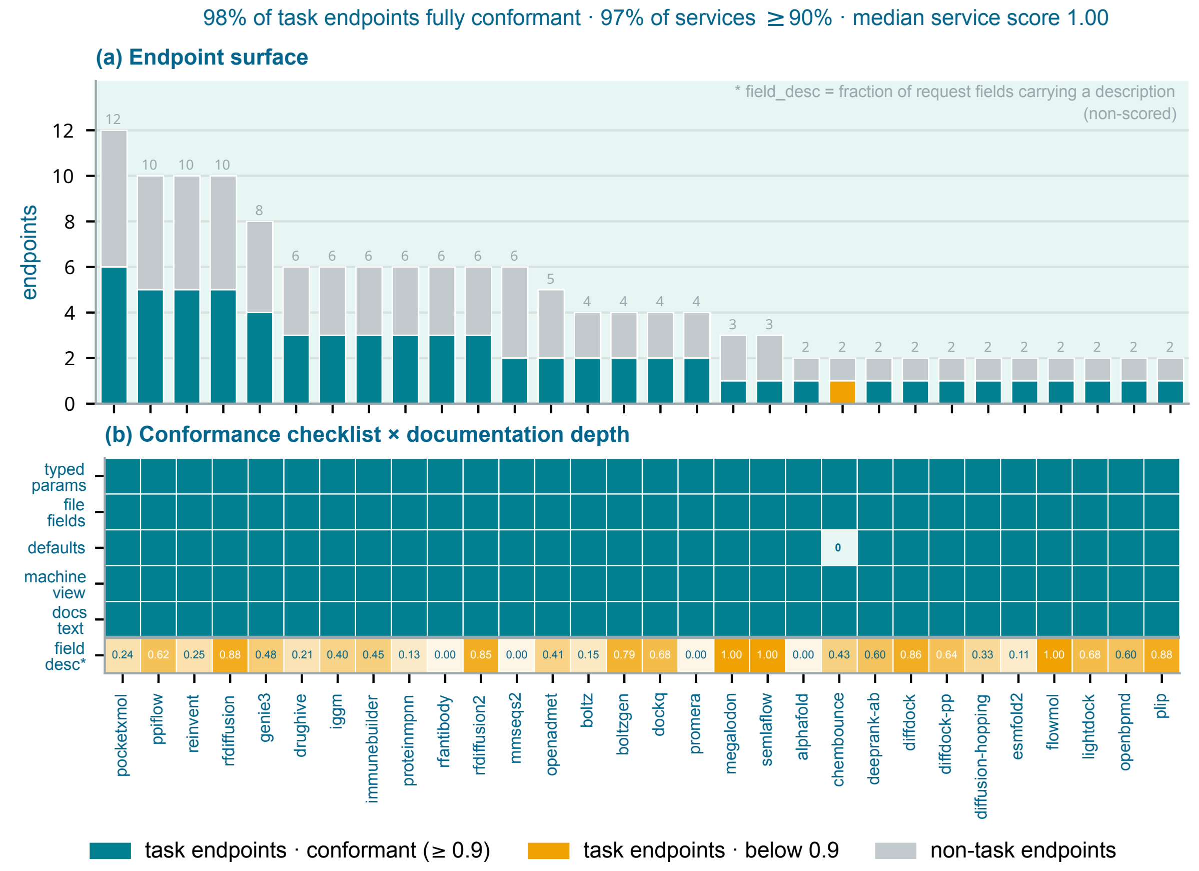


**Figure S1:** Fleet-wide audit of the uniform, self-describing contract. A manifest-only audit of the 30 deployed services (141 endpoints, 68 of them `bioq run` task endpoints) scores every endpoint against a five-item checklist — the structural items `typed_params`, `file_fields`, and `machine_view`, and the metadata items `defaults` and `docs_text`. 98.5 % of task endpoints are fully conformant and 96.7 % of services sit at or above a 0.9 score (median 1.0); the single residual gap is one unannotated `input_smiles_uri` default on `chembounce scaffold_hop`, which falls below the bar (amber). (a) Endpoint surface, per service. Each vertical bar is a service's total endpoint count — a dark base for task (async job interface) endpoints and a light top for non-task (sync job and status interface) endpoints. Teal marks services at or above the 0.9 bar; amber marks those below it. (b) Conformance checklist × documentation depth. One column per service and one row per contract item. The five scored rows report the fraction of task endpoints passing each checklist item; the final `field_desc` row (non-scored) reports the mean fraction of request fields carrying a description.

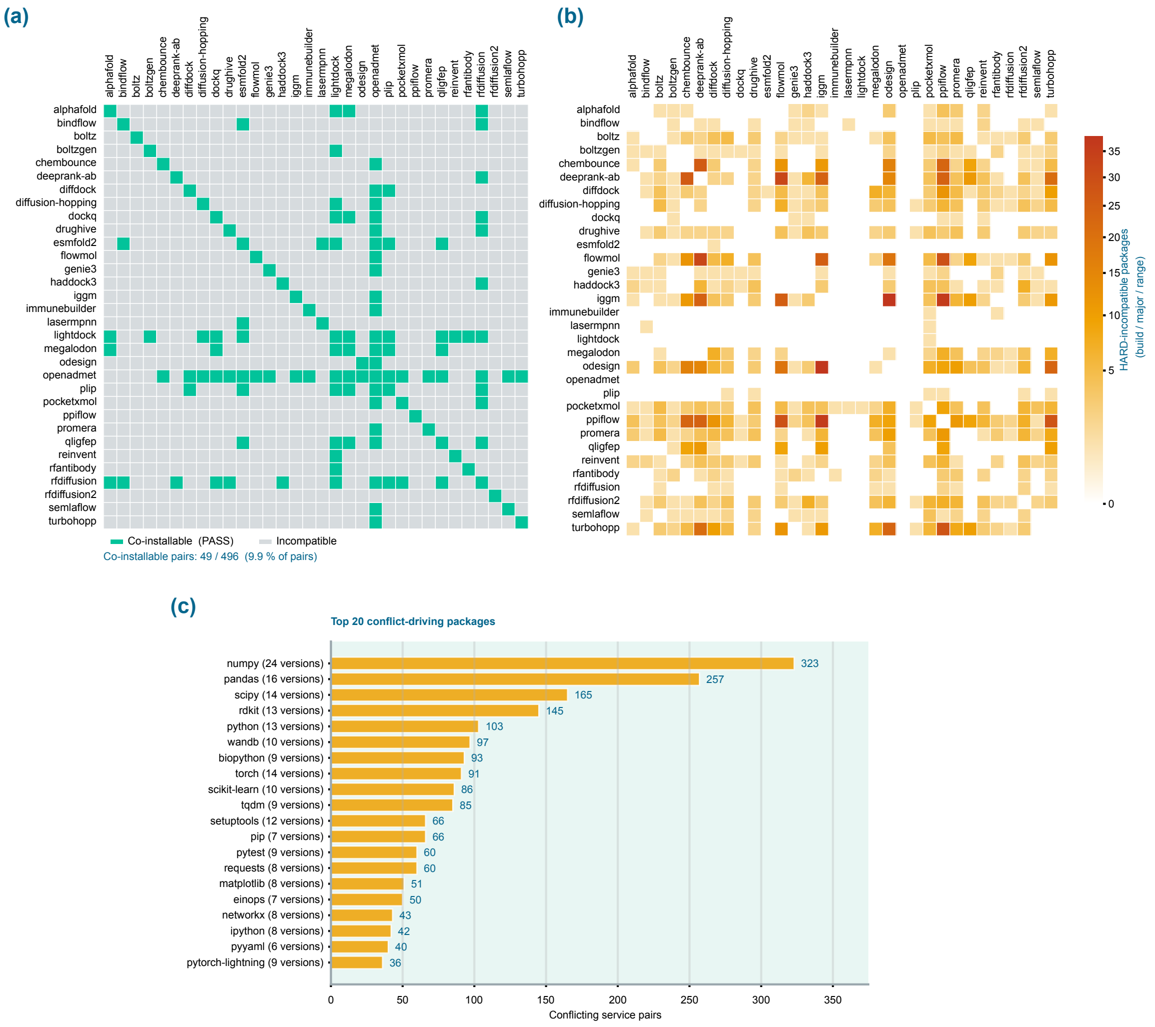


**Figure S2:** Declared-dependency incompatibility of the bioq fleet (the 32 tools with parseable upstream dependency specifications; 496 pairs). (a) Co-installability: whether two tools' declared dependencies can literally share one environment as-is (green = co-installable, grey = not; 49 of 496 pairs, 9.9 %). (b) HARD-conflict heatmap: disjoint version ranges per tool pair — no single version satisfies both. (c) Top 20 conflict-driving packages.

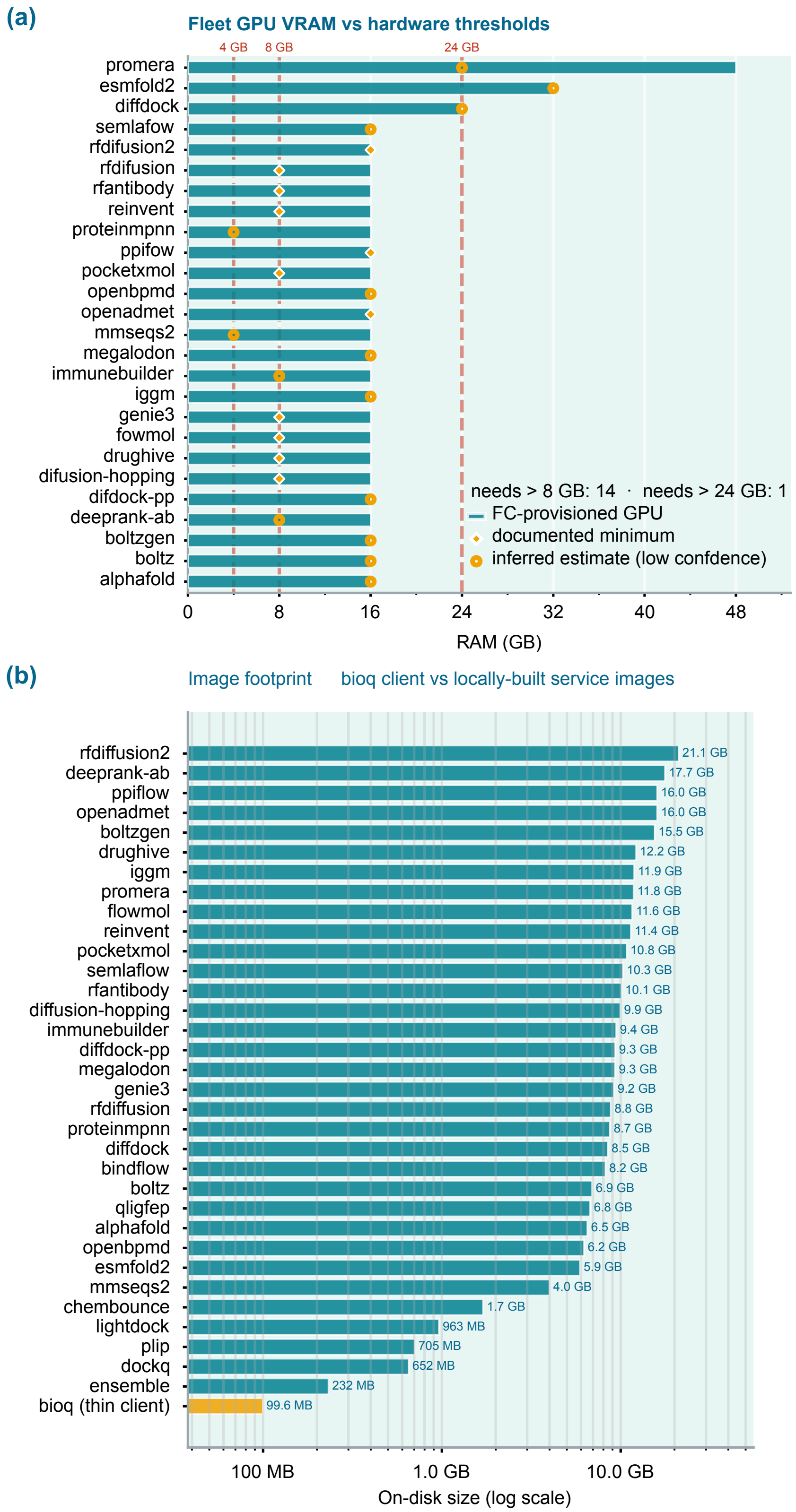


**Figure S3:** Compute barrier for the full services fleet. (a) GPU VRAM consumption per service. VRAM requirements marked as inferred are best-known estimates (15 of 26). The fleet cannot run on laptop GPUs. (b) Storage footprint of the bioq client and service Docker images (model weights excluded from Docker images).

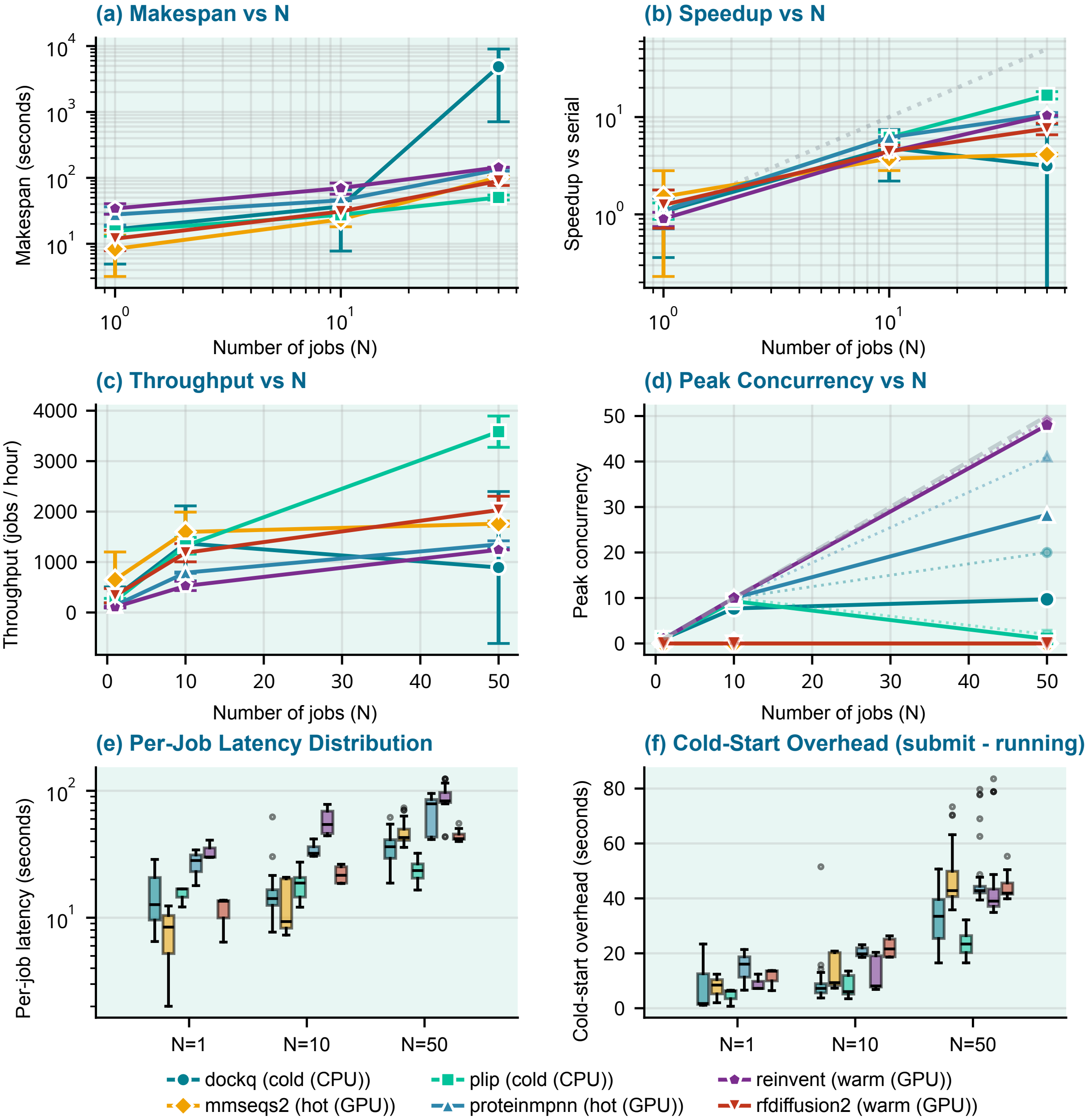


**Figure S4:** Throughput scaling of bioq serverless fan-out on Alibaba Cloud Function Compute (FC). Six representative services spanning three serverless tiers — cold CPU (`dockq`, `plip`), hot GPU (`mmseqs2`, `proteinmpnn`), and warm GPU (`reinvent`, `rfdiffusion2`) — are measured as a function of batch size $N \in \{1, 10, 50\}$; the upper bound $N = 50$ is the FC GPU instance-quota ceiling. Each point is the mean over three independent replicates, with error bars showing $\pm 1$ standard deviation. Colour and marker identify the service (shared legend beneath the panels). (a) Makespan vs $N$ (log–log), wall-clock to finish all N jobs; (b) Speedup vs $N$ (log–log), $N\times$ single-job time / makespan, relative to the serial baseline; (c) Throughput vs $N$, completed jobs / makespan; (d) Peak concurrency vs $N$, max simultaneous worker instances observed; (e) Per-job latency distribution (log-scale box plots), wall-clock time for a single job from submit to completion; (f) Cold-start overhead (box plots), time from submit to first running status.

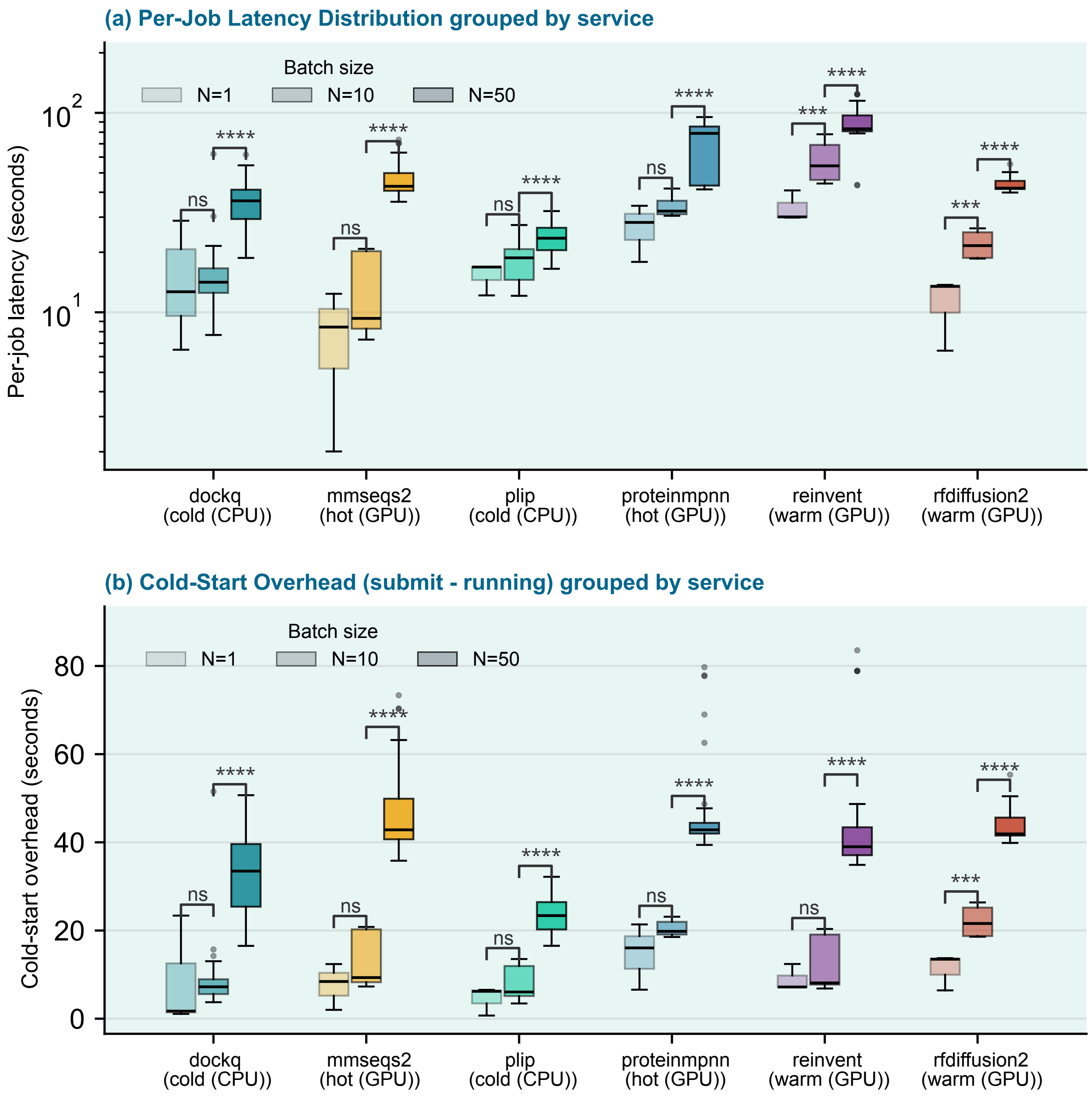


**Figure S5:** Per-job latency and cold-start overhead, grouped by service, on FC. The x-axis lists six services; within each service, three box plots show batch size $N = 1, 10, 50$ (lighter → darker). Boxes pool all jobs across three replicates. Brackets show two-sided Mann–Whitney U tests between adjacent $N$ levels. (a) Per-job latency (seconds, y-axis log scale); (b) Cold-start overhead (seconds).

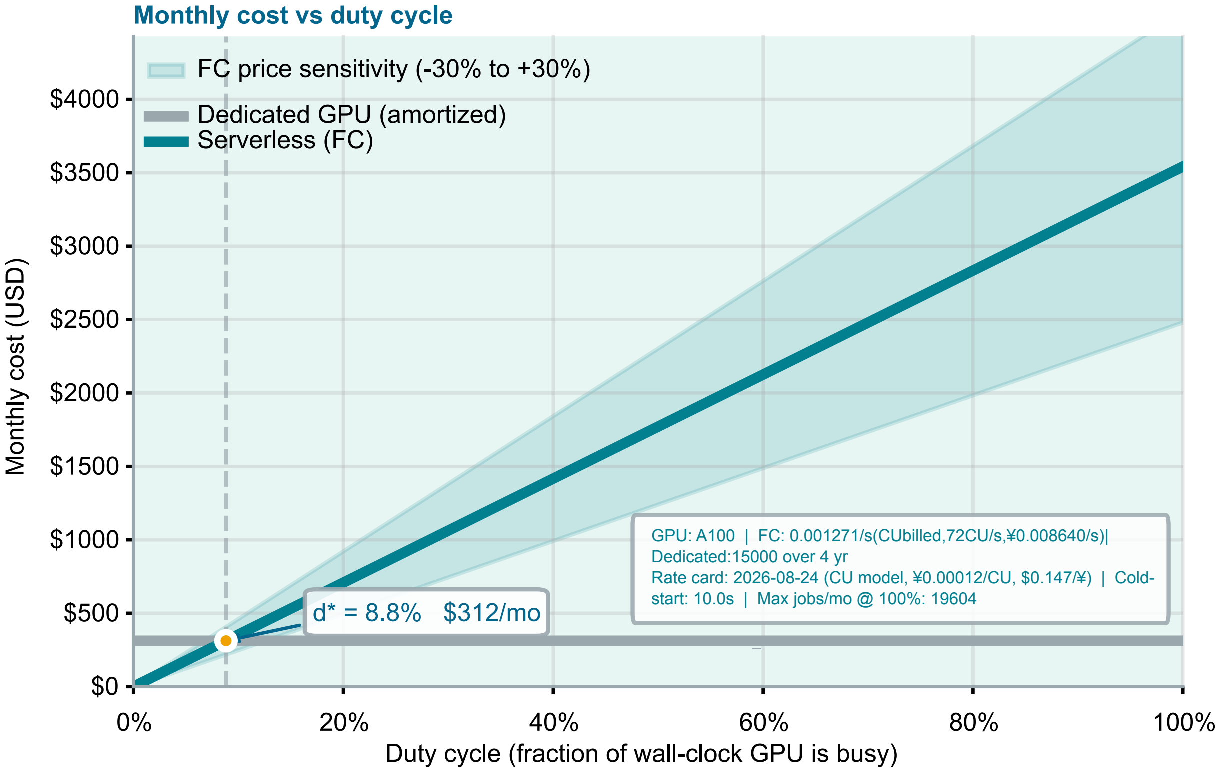


**Figure S6:** Monthly cost and duty-cycle break-even for serverless and dedicated A100-class GPU capacity. The bioq serverless estimate uses the Alibaba Cloud Function Compute pricing schedule (2026). Under the first pricing tier, a 40-GB Ampere-class allocation consumes 72 $\mathrm{CU\,s^{-1}}$ at CNY 0.00012 per CU, equivalent to USD 4.57 $\mathrm{h^{-1}}$ at USD 1 = CNY 6.80. The calculation includes 10 s of billed cold-start overhead per invocation and yields a weighted mean cost of approximately USD 0.18 per observed job. The owned-A100 comparison assumes a purchase price of USD 15,000 amortized over four years (USD 0.43 $\mathrm{h^{-1}}$), whereas the persistent cloud-VM comparison assumes USD 2.50 $\mathrm{h^{-1}}$. The resulting break-even duty cycles are approximately 8.8 % and 52 %, respectively; below each crossover, serverless execution is less expensive. The shaded band shows the price-sensitivity sweep. The ownership estimate excludes host-system, electricity, cooling, maintenance, financing, storage, and network-transfer costs and is therefore conservative in favor of owned hardware.

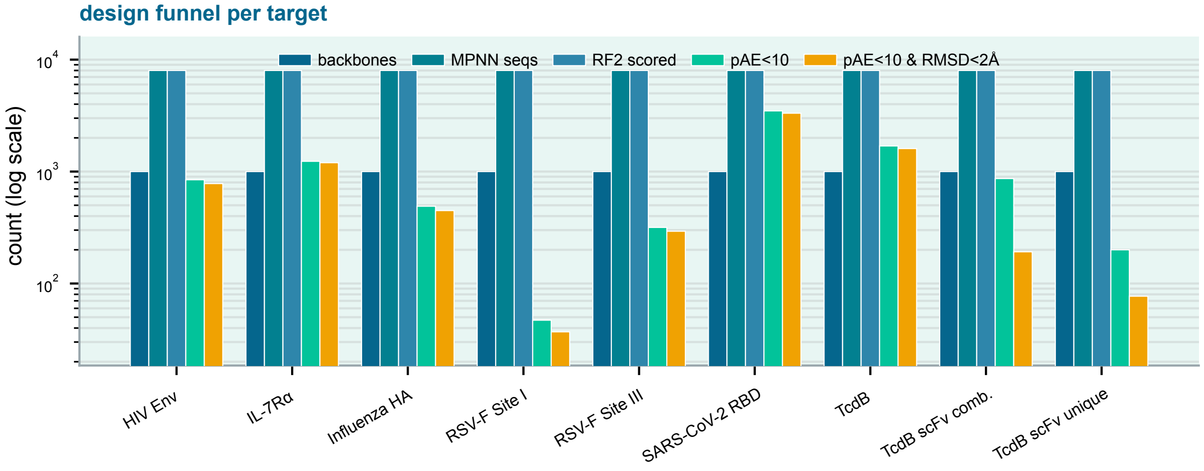


**Figure S7:** Per-target design funnel for the nine-target RFantibody de novo antibody-design campaign run entirely through bioq (three `bioq run` calls per target, no local GPU). For each target, five grouped vertical bars report the count at successive pipeline stages: RFdiffusion backbones (1,000), ProteinMPNN sequences (8,000 = 8 sequences per backbone), sequences carried through RF2 structure prediction and scoring (8,000), designs passing the interface-pAE $< 10$ filter, and designs passing the combined acceptance criterion (interface pAE $< 10$ *and* design–model CDR RMSD $< 2$ Å, self-consistency RMSD, design vs RF2-predicted structure; amber). The y-axis is log-scaled.

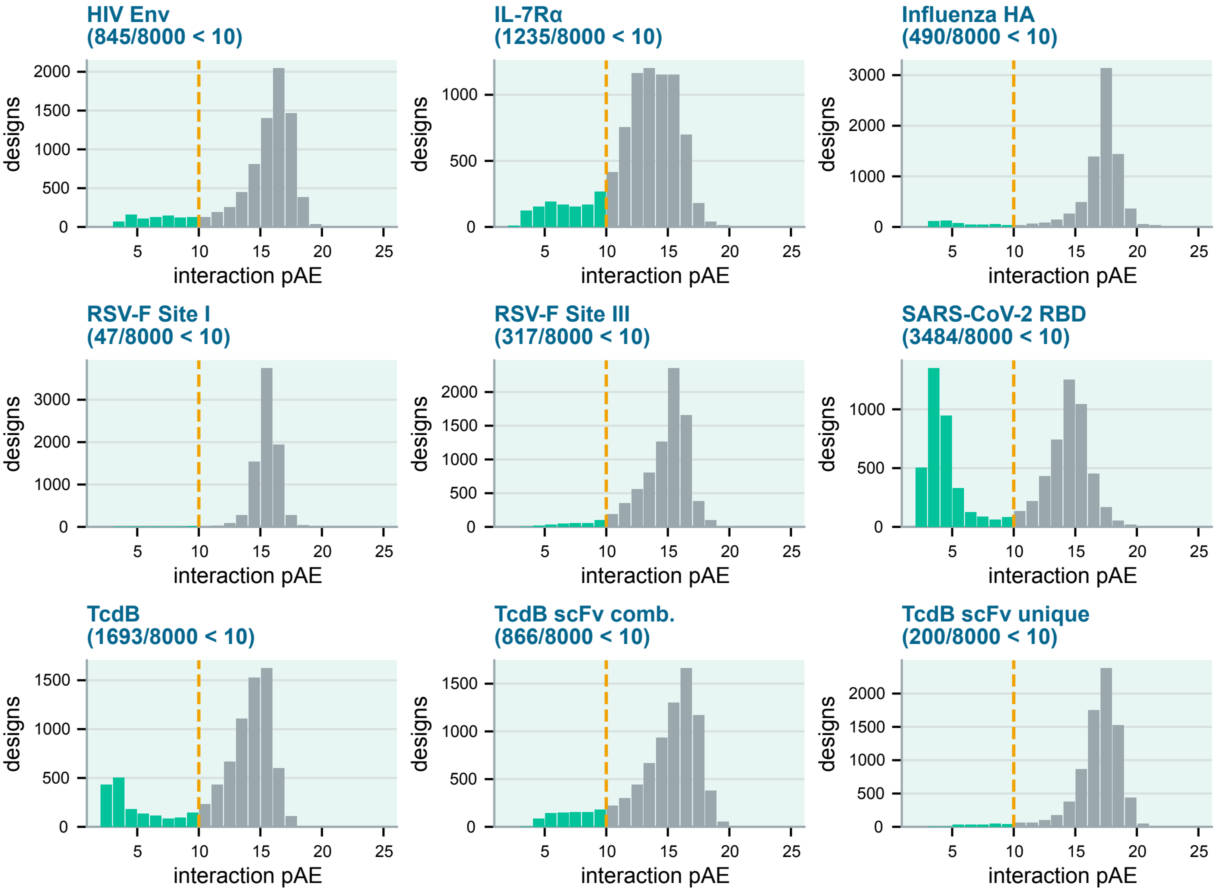


**Figure S8:** Distribution of interface pAE (`interaction_pae`, a.k.a. `pae_interaction` in the paper) across all 8,000 designs per target, one histogram per target in a 3×3 grid. Green bars mark designs passing the pAE $< 10$ filter, grey bars the failures, and the dashed amber line the threshold. All panels share a single binning (2–26 in unit steps); each panel title reports the pass count over the total.

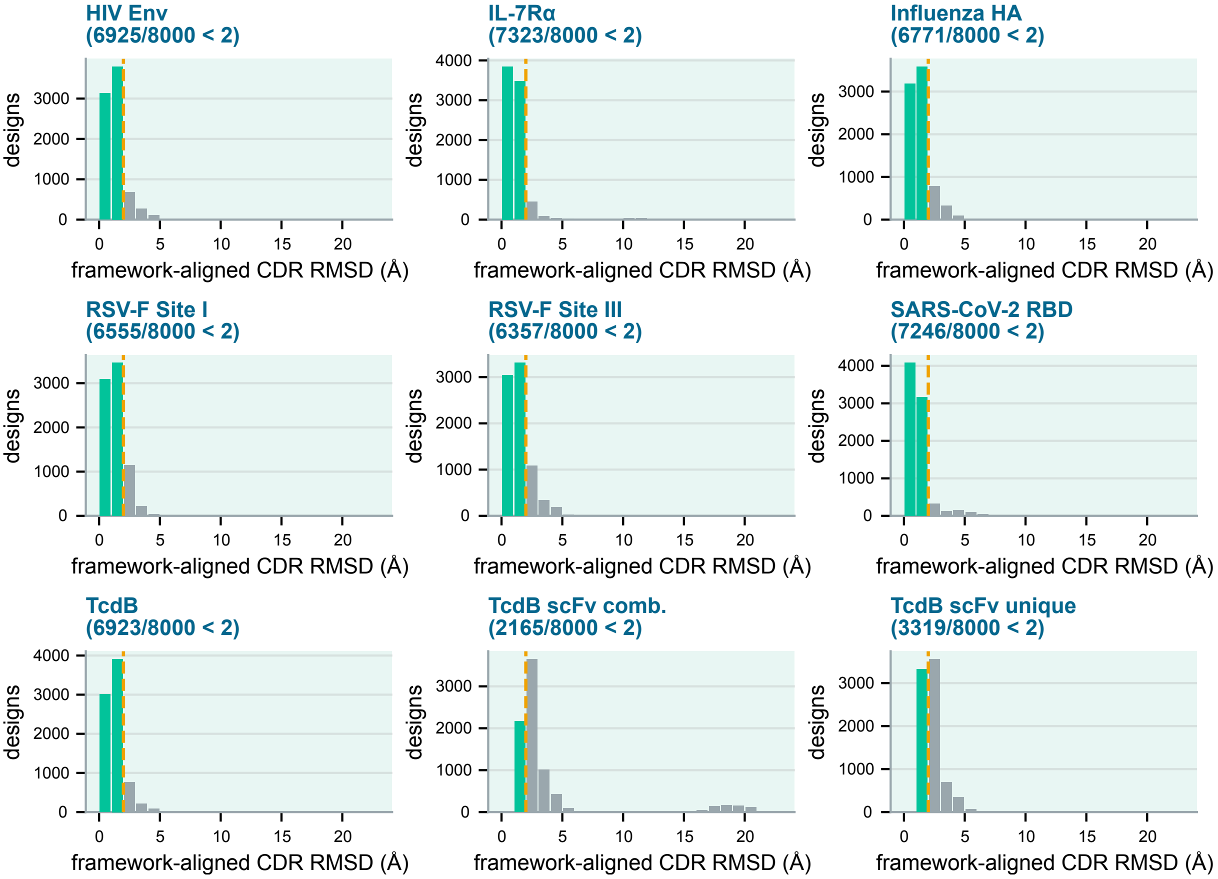


**Figure S9:** Distribution of design–model CDR RMSD across all 8,000 designs per target, one histogram per target in a 3×3 grid. Green bars mark designs passing the RMSD < 2 Å filter, grey bars the failures, and the dashed amber line the 2 Å threshold. All panels share a single binning (0–24 Å in 1 Å steps).

### Supplementary Tables

**Table S1:** The 38-service bioq fleet. Short names omit the `-server` suffix. Stages: 1 Structure prediction, 2 Sequence search, 3 De novo design, 4 Docking & screening, 5 Affinity / free energy, 6 ADMET / PK, 7 Scoring & QC. Modalities: protein, antibody/VHH, small molecule, peptide, nucleic acid, cross-modal. Citations are author–year.

| **Short name** | **Stage** | **Modality** | **Upstream method (citation)** |
|---|---|---|---|
| `alphafold` | Structure prediction | protein / cross-modal | AlphaFold / AlphaFold-Multimer (Jumper et al., 2021; Abramson et al., 2024; Evans et al., 2021) |
| `boltz` | Structure prediction | cross-modal | Boltz-2 (Passaro et al., 2025; Wohlwend et al., 2024) |
| `esmfold2` | Structure prediction | protein / antibody/VHH | ESMFold2 (Lin et al., 2023) |
| `immunebuilder` | Structure prediction | antibody/VHH | ImmuneBuilder (Abanades et al., 2023) |
| `promera` | Structure prediction | protein / antibody/VHH | Promera (Jing et al., 2026) |
| `ensemble` | Structure prediction | antibody/VHH / protein | Multi-method ensemble (AlphaFold / ESMFold / Boltz / Promera) |
| `mmseqs2` | Sequence search | protein / cross-modal | MMseqs2 (Steinegger and Söding, 2017) |
| `diamond` | Sequence search | protein / cross-modal | DIAMOND (Buchfink et al., 2026) |
| `rfdiffusion` | De novo design | protein | RFdiffusion (Watson et al., 2023) |
| `rfdiffusion2` | De novo design | protein | RFdiffusion2 (Ahern et al., 2026; Krishna et al., 2024) |
| `genie3` | De novo design | protein | Genie 3 (Lin et al., 2026) |
| `ppiflow` | De novo design | protein / antibody/VHH | PPIFlow (Yu et al., 2026) |
| `boltzgen` | De novo design | protein / antibody/VHH / peptide | BoltzGen (Stark et al., 2025) |
| `rfantibody` | De novo design | antibody/VHH | RFantibody (RFdiffusion + ProteinMPNN + RoseTTAFold-2) (Bennett et al., 2026; Watson et al., 2023; Dauparas et al., 2022; Baek et al., 2021) |
| `iggm` | De novo design | antibody/VHH | IgGM (Wang et al., 2025) |
| `odesign` | De novo design | protein / nucleic acid / small molecule | ODesign (Zhang et al., 2025) |
| `proteinmpnn` | De novo design | protein | ProteinMPNN (Dauparas et al., 2022) |
| `lasermpnn` | De novo design | protein / small molecule | LASErMPNN (+ LigandMPNN) (Fry et al., 2026; Dauparas et al., 2022) |
| `flowmol` | De novo design | small molecule | FlowMol3 (Dunn and Koes, 2025) |
| `semlaflow` | De novo design | small molecule | SemlaFlow (Irwin et al., 2025) |
| `megalodon` | De novo design | small molecule | Megalodon (Reidenbach et al., 2025) |
| `reinvent` | De novo design | small molecule | REINVENT (Loeffler et al., 2024; Blaschke et al., 2020) |
| `pocketxmol` | De novo design | small molecule | PocketXMol (Peng et al., 2026) |
| `drughive` | De novo design | small molecule | DrugHIVE (Weller and Rohs, 2024) |
| `chembounce` | De novo design | small molecule | ChemBounce (Jang et al., 2025) |
| `diffusion-hopping` | De novo design | small molecule | DiffHopp (Torge et al., 2023) |
| `turbohopp` | De novo design | small molecule | TurboHopp (Yoo et al., 2024) |
| `diffdock` | Docking & screening | small molecule | DiffDock (Corso et al., 2023) |
| `diffdock-pp` | Docking & screening | protein | DiffDock-PP (Ketata et al., 2023) |
| `lightdock` | Docking & screening | protein | LightDock (Jiménez-García et al., 2018) |

*continued on next page*

*Table S1 (continued)*

| Short name | Stage | Modality | Upstream method (citation) |
|---|---|---|---|
| `haddock3` | Docking & screening | protein / cross-modal | HADDOCK3 (Giulini et al., 2025) |
| `plip` | Docking & screening | small molecule / protein | PLIP (Schake et al., 2025) |
| `bindflow` | Affinity / free energy | small molecule | BindFlow (León et al., 2026) |
| `qligfep` | Affinity / free energy | small molecule | QligFEP (Alencar Araripe et al., 2025) |
| `openbpmd` | Affinity / free energy | small molecule | OpenBPMD (Lukauskis et al., 2022) |
| `openadmet` | ADMET / PK | small molecule | OpenADMET (Fraser et al., 2026) |
| `dockq` | Scoring & QC | protein | DockQ (Basu and Wallner, 2016; Mirabello and Wallner, 2024) |
| `deeprank-ab` | Scoring & QC | antibody/VHH | DeepRank-Ab (Xu et al., 2025) |

**Table S2:** Per-target in-silico design funnel for the nine-target RFantibody de novo antibody-design campaign (seven VHH, two scFv), run entirely through bioq. Columns, left to right: RFdiffusion backbones; ProteinMPNN sequences (eight per backbone); sequences carried through RF2 folding and scoring; designs passing the interface-pAE $< 10$ filter; and designs passing the combined acceptance filter (interface pAE $< 10$ *and* design–model CDR RMSD $< 2$ Å). The final column reports the published RFantibody per-target in-silico success rate for comparison. Targets are ordered by the best-of-8 backbone pass rate.

| Target | Type | Backbones | Sequences | Scored | pAE <10 | Passed | Passed (%)$^\dagger$ | Best-of-8 (%)$^\ddagger$ | Published (%)$^\S$ |
|---|---|---|---|---|---|---|---|---|---|
| SARS-CoV-2 RBD | VHH | 1,000 | 8,000 | 8,000 | 3,484 | 3,321 | 41.51 | 70.6 | 28.0 |
| TcdB | VHH | 1,000 | 8,000 | 8,000 | 1,693 | 1,605 | 20.06 | 48.9 | 41.0 |
| IL-7R$\alpha$ | VHH | 1,000 | 8,000 | 8,000 | 1,235 | 1,200 | 15.00 | 43.0 | 34.5 |
| HIV Env | VHH | 1,000 | 8,000 | 8,000 | 845 | 781 | 9.76 | 34.2 | – |
| RSV-F Site III | VHH | 1,000 | 8,000 | 8,000 | 317 | 293 | 3.66 | 17.2 | 25.0 |
| Influenza HA | VHH | 1,000 | 8,000 | 8,000 | 490 | 448 | 5.60 | 14.6 | 11.5 |
| RSV-F Site I | VHH | 1,000 | 8,000 | 8,000 | 47 | 37 | 0.46 | 2.4 | – |
| TcdB scFv (comb.) | scFv | 1,000 | 8,000 | 8,000 | 866 | 192 | 2.40 | 9.6 | – |
| TcdB scFv (unique) | scFv | 1,000 | 8,000 | 8,000 | 200 | 77 | 0.96 | 5.8 | – |

$^\dagger$ Fraction of scored sequences passing the combined filter. $^\ddagger$ Fraction of backbones whose best-scoring of eight MPNN sequences passes the combined filter; this per-backbone best-of-8 rate is the metric reported by the published RFantibody method and gives the main-text pass-rate range. $^\S$ Published per-target in-silico success rate of the RFantibody method (Bennett et al., 2026); a dash marks targets with no reported per-target screening.
